# Amelioration of DSS-induced Acute Colitis in Mice by Recombinant Monomeric Human Interleukin-22

**DOI:** 10.1101/2021.11.04.467038

**Authors:** Suhyun Kim, Eun-Hye Hong, Cheol-Ki Lee, Yiseul Ryu, Hyunjin Jeong, Seungnyeong Heo, Joong-jae Lee, Hyun-Jeong Ko

**Author notes:** Correspondence should be addressed to J.-j.L. and H.-J.K. These authors contributed equally to this work (Suhyun Kim, Eun-Hye Hong, Cheol-Ki Lee).

## Abstract

Interleukin-22 (IL-22), a pleiotropic cytokine, is known to have a profound effect on the regeneration of damaged intestinal barriers. The tissue-protective properties of IL-22 are expected to be potentially exploited in the attenuation and treatment of colitis. However, because of the disease-promoting role of IL-22 in chronic inflammation, a comprehensive evaluation is required to translate IL-22 into the clinical domain. Here, we present the effective production of soluble human IL-22 in bacteria to prove whether recombinant IL-22 has the ability to ameliorate colitis and inflammation. IL-22 was expressed in the form of a biologically active monomer and a non-functional dimer. Monomeric IL-22 (mIL-22) was highly purified through a series of three separate chromatographic methods and an enzymatic reaction. We reveal that the resulting mIL-22 is correctly folded and is able to phosphorylate signal transducer and activator of transcription 3 in HT-29 cells. Subsequently, we demonstrate that mIL-22 enables the attenuation of dextran sodium sulfate-induced acute colitis in mice, as well as the suppression of pro-inflammatory cytokine production. Collectively, our results suggest that the recombinant mIL-22 is suitable to study the biological roles of endogenous IL-22 in immune responses and can be developed as a biological agent associated with inflammatory disorders.

## Introduction

Interleukin-22 (IL-22), a member of the IL-10 cytokine family, plays a significant role in host defense against microbial infections and a broad range of immunological disorders including psoriasis, Crohn’s disease, hepatitis, and rheumatoid arthritis.[1] IL-22 is characterized by a six alpha helix bundle with two disulfide bonds[2, 3], which mainly produced in various immune cells such as T helper cells (T_H_1, T_H_17, and T_H_22), type 3 innate lymphoid cells, natural killer T cells, and γδ T cells.[4-6] Interestingly, contrary to the original meaning of interleukin, IL-22 can mediate cellular communication between immune cells and some non-immune cells to induce the expression of tissue-specific effector molecules for cell differentiation and tissue regeneration.[7, 8] The biological functions of IL-22 are triggered by formation of binary complex between IL-22 and IL-22 receptor 1 (IL-22R1), and the dimer is subsequently interacted with IL-10 receptor 2 (IL-10R2).[9-11] The resulting complexes can activate downstream signaling cascades by phosphorylation of Janus kinase 1 (JAK1) and signal transducer and activator of transcription 3 (STAT3), giving rise to upregulation of many relevant genes.[12, 13] IL-22R1 has high-binding affinity toward IL-22 and is restrictedly expressed on epithelial cells unlike IL-10R2 expressed ubiquitously.[7] Therefore, the major target organs of IL-22 are generally considered as liver, pancreas, lung, colon, intestine and kidney in which epithelial cells cooperatively form a continuous physical barrier to allow tightly regulated cellular transport and prevent pathogen infections.[3]

As pleiotropic functions of IL-22 have been elucidated, many studies have demonstrated the therapeutic potential of IL-22 in liver and pancreas regeneration, infection prevention and graft-versus-host disease.[5, 14] Recently, a recombinant human IL-22-Fc dimer (F-562) was evaluated in phase II clinical trials to provide evidence whether IL-22 could be a promising agent to treat patients with alcoholic hepatitis.[14-17] This study clearly showed that the engineered IL-22 was resulted in improved liver injury-related scores in patients by showing both decreased plasma inflammatory markers and elevated regenerative markers.[15] More importantly, IL-22 seems to be critical in inflammatory bowel disease (IBD) such as Crohn’s disease and ulcerative colitis.[1, 13, 18, 19] Several studies provided the protective roles of IL-22 in IBD by promoting regeneration of intestinal stem cells (ISCs) and attenuating dysregulated immune responses. As above mentioned, IL-22 can selectively activate the STAT3 pathway, thereby upregulating the expression of several target genes primarily involved in cell survival and proliferation. Consequently, in damaged intestinal tissue, IL-22 substantially promotes the re-establishment of the intestinal epithelial barrier integrity and function.[1] Based on the molecular mechanism, it was suggested that IL-22 is essential for epithelial maintenance and regeneration in intestinal injury.[20] As the beneficial effects of IL-22 have been revealed, IL-22-mediated STAT3 activation is attracting attention as a promising therapeutic option for intestinal wound healing and inflammatory disorders such as necrotizing enterocolitis (NEC).[18, 21, 22]

Despite the clinical significance of IL-22, the effective production of human recombinant IL-22 (hrIL-22) in bacteria has not yet been established. Unfortunately, rIL-22 has been mainly obtained by a refolding method from bacterial inclusion bodies due to its extremely low solubility.[23, 24] Moreover, because the renaturation process is a complex and labor-intensive procedure, it is difficult to optimize and perform at laboratory scale for research purposes. In this study, we present a cost-effective production of soluble hrIL-22 in bacteria without a solubilization step, and evaluate its biological activities through *in vitro* cell-based assay and *in vivo* mouse studies. Briefly, to promote the correct formation of two disulfide bonds of IL-22, we genetically inserted a thioredoxin (Trx) tag at the N-terminus of IL-22 and the resulting IL-22 exhibited highly enhanced soluble expression.[25] Through a series of downstream processing, monomeric IL-22 (mIL-22) was finally purified with very high purity (up to 95%). We subsequently characterized the structural and biological properties of mIL-22 by several biochemical studies, including circular dichroism analysis and cell proliferation assay. Finally, a therapeutic potential of recombinant IL-22 was successfully demonstrated in a dextran sodium sulfate (DSS)-induced acute colitis mouse model.[18, 21, 26] Details are reported herein.

## Materials and Methods

### Molecular Cloning

Codon optimized human interleukin-22 (IL-22) gene for *Escherichia coli* was synthesized (IDTdna, USA) and amplified through polymerase chain reaction (PCR) using the following primers (Restriction enzyme sites are underlined): 1) forward 5’- ATATAT<u>CCATGG</u>CTCCCATTTCGAGTCATTGTCG-3’ 2) reverse 5’- ATATAT<u>CTCGAG</u>GATACAAGCGTTGCGTAAGG-3’. Forward and reverse primers bear *NcoI* and *XhoI* cleavage sites, respectively. For tagging thioredoxin (Trx) at the N-terminal end of IL-22, the amplified IL-22 gene was cloned into a pET32a (+) vector (Novagen, USA) using two restriction sites as shown in **Figure 1b**. The final construct of recombinant IL-22 possessed both a Trx tag and a 6x histidine tag, allowing for increased soluble expression and facilitated affinity purification.

**Figure 1.**
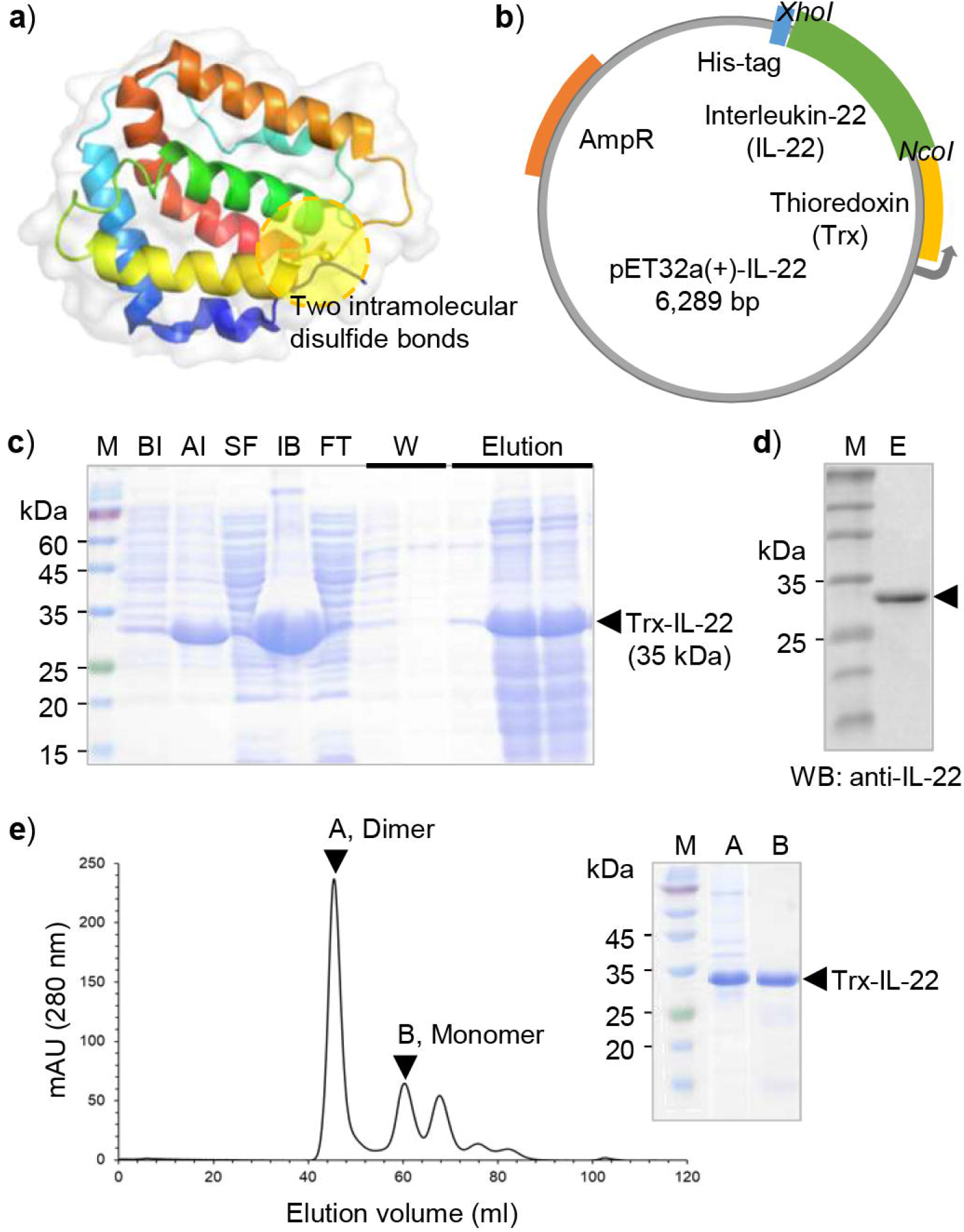
Bacterial expression and purification of thioredoxin-tagged human interleukin-22. **a)** The ribbon representation of the folded structure of interleukin-22 (IL-22). Two intramolecular disulfide bonds were indicated within yellow transparent circle. **b)** A schematic diagram of a designed bacterial expression vector pET32a (+) for the production of IL-22 with thioredoxin (Trx). **c)** SDS-PAGE analysis of Trx-IL-22 protein which purified by using nickel-nitrilotriacetic acid (Ni-NTA) resin. BI: Before IPTG induction. AI: After IPTG induction. SF: Soluble fraction. IB: Inclusion body. FT: Flow through. W: Washing fraction. **d)** Western blot analysis of the purified Trx-IL-22 in elution fraction shown in Figure 1c using anti-IL-22 primary antibody. **e)** Size-exclusion chromatography profile of Trx-IL-22 protein. Peaks A and B containing dimer and monomer Trx-IL-22, respectively, were identified by SDS-PAGE analysis.

### Protein Expression and Purification

The expression plasmid vector pET32a (+) containing thioredoxin fused human interleukin-22 (Trx-IL-22) gene was transformed into Origami B (DE3) competent cells, then the transformed cells were incubated on LB broth agar plates containing ampicillin (100 μg/ml), kanamycin (50 μg/ml), and tetracycline (10 μg/ml) at 37°C overnight. A single colony was picked, inoculated into 10 ml of LB broth medium, and pre-cultured at 37°C overnight. Pre-cultured cells were diluted 100-fold in 1000 ml of LB medium with all three types of antibiotics. When the optical density at 600 nm (OD_600_) of the cultured cells reached 0.5-0.7, Isopropyl-ß-D- thiogalactopyranoside (IPTG) was added at final concentration of 0.1 mM to induce protein expression. The ideal induction time in this study was approximately 21 hrs at 18°C. Cultured cells were harvested by centrifugation at 6000 rpm for 10 min at 4°C. The pelleted cells of 1000 ml culture were re-suspended with 40 ml of lysis buffer (50 mM NaH_2_PO_4_ (pH 8.0), 300 mM NaCl, and 10 mM imidazole).

Prior to cell disruption, protease inhibitor cocktail (Roche, USA) was added, and sonication was conducted at 40% amplitude for 1 hour. After sonication, cell lysate was separated by centrifugation at 13,000 rpm for 1 hour at 4°C. Inclusion bodies were solubilized in 8 M urea for SDS-PAGE analysis. The soluble fraction was filtered with a syringe filter with 0.22 μm pore size, and further purified by affinity chromatography. First, Ni-NTA agarose resin (Qiagen, Germany) was equilibrated in a disposable 5 ml polypropylene column (Thermo, USA) with lysis buffer for 10 column volume. The filtered solution was applied to the buffer-equilibrated Ni-NTA column. The column was washed three times with 5 column volume of a wash buffer (50 mM NaH_2_PO_4_ (pH 8.0), 300 mM NaCl, and 20 mM imidazole). Bound proteins were eluted by elution buffer (50 mM NaH_2_PO_4_ (pH 8.0), 300 mM NaCl, and 250 mM imidazole). Subsequently, size exclusion chromatography using HiLoad® 16/600 Superdex® 75 column (Cytiva, USA) was performed to increase purity of Trx-IL-22. Overall purity and concentration of the purified protein were determined by SDS-PAGE and Bradford assay, respectively.

### Thioredoxin Tag Removal

To remove thioredoxin (Trx) tag from interleukin-22 (IL-22) protein, enterokinase was employed to cleave its recognition site located between Trx and IL-22. Buffer exchange of purified Trx-IL-22 was performed with an enterokinase reaction buffer (20 mM Tris-HCl (pH 8.0), 50 mM NaCl and 2 mM CaCl_2_) using a PD-10 desalting column (GE Healthcare, USA). Light chain of human enterokinase (10 μg/ml) was incubated with Trx-IL-22 (0.4 mg/ml) for 2 hrs at 25°C. Following enzymatic reaction, protease inhibitor cocktail (Roche, USA) was treated to the Trx-IL-22 solution to prevent unwanted non-specific protein cleavage and degradation. Buffer exchange was conducted with A buffer (20 mM Tris, pH 7.8), and anion exchange chromatography was sequentially carried out to isolate IL-22 from the enterokinase treated solutions. IL-22 and other proteins were separated through a HiTrap Q FF column (Cytiva, USA) under a linear salt gradient condition from 0% to 50% B buffer (20 mM Tris, 1 M NaCl, and pH 7.8). Respective eluted peaks were analyzed by SDS-PAGE, and all fractions containing IL-22 were concentrated using a 10 kDa centrifugal filter (Merck Millipore, USA).

### *In vivo* mouse models

C57BL/6 background female 8-week-old mice were purchased from Koatech (Pyeongtaek, korea). All experiments were performed by the Institutional Animal Care and Use committee of Kangwon University (IACUC admission number KW-200831-1) and the mice were bred during experiment in the Animal Laboratory Center of Kangwon National University. For acute colitis induction, mice were treated with 2.5 % dextran sulfate sodium (wt/vol, molecular weight 36-50 kDa; MP Biomedicals, Solon, OH, USA) in drinking water for 5 consecutive days with 4 additional days of tap water only. We assessed the induction of colitis through body weight changes. Colon length was assessed by millimeter ruler on the 9 day of the experiment.

### Enzyme-linked Immunosorbent Assay

Cytokines and chemokines including mouse IL-1β, IL-6, IL-17A, CXCL1, and CCL2 were measured using Mouse Uncoated ELISA Kit (Invitrogen, Vienna, Austria) and mouse CXCL1/KC DuoSet ELISA (R&D systems, Minneapolis, MN, USA). Briefly, mouse colon tissues were homogenized using Minilys personal homogenizer (Bertin, Montigny-le-Bretonneux, France). Tissue homogenate was centrifuged and used only supernatant. Immuno-plates (Thermo Fisher scientific, Waltham, MA, USA) were coated with capture antibody at 4°C for overnight and Ab-coated wells were blocked with 1x assay diluent for 1 hr at 20°C. Standard and sample were diluted and put into each well at 4°C for 18 h. The detection Abs conjugated with HRP were applied into wells and wells were developed with 1x TMB substrate. The absorbance was measured at 450 nm by using SpectraMax i3(Molecular Devices, San Jose, CA, USA).

### Circular Dichroism

Circular dichroism (CD) spectrum of interleukin-22 (IL-22) was measured at 25°C with J-1500 CD spectropolarimeter (Jasco, Japan) using a 1 mm path cuvette. IL-22 (0.5 mg/ml) was dissolved in a tris-based buffer (20 mM Tris (pH 7.8) and 75 mM NaCl). The collected CD data were analyzed by a K2D3 server (http://cbdm-01.zdv.uni-mainz.de/~andrade/k2d3) to estimate the secondary structure of IL-22.

### Western Blot

Purification of recombinant interleukin-22 (IL-22) was demonstrated through western blot analysis. Thioredoxin-tagged IL-22 and tag-removed IL-22 were loaded onto 15% SDS-PAGE, transferred to a polyvinylidene fluoride (PVDF) membrane. After masking the membrane with a blocking buffer (Tris-buffered Saline (TBS) containing 0.1% Tween-20 and 2% BSA) for 30 min, the membrane was incubated with mouse anti-human IL-22 antibody (1 μg/ml; R&D systems, USA) overnight at 4°C. The membrane was washed three-times with TBST (TBS containing 0.1% Tween-20) and incubated with HRP-conjugated mouse IgG kappa binding protein (1:5000; Santa Cruz Biotechnology, USA) for 2 hrs at room temperature. After washing the membranes with TBST, protein bands were detected with a chemiluminescent substrate (BIOMAX, USA) through a CCD Imager (GE life sciences, USA).

To verify IL-22 induced signal transducer and activator of transcription 3 (STAT3) phosphorylation, human colon cancer HT-29 cells (ATCC, USA) in RPMI 1640 media (Welgene, Korea) were seeded at 1 × 10^6^ cells/well on a six-well plate. After 24 hours, 10, 100 and 1000 ng/ml of IL-22 diluted in serum-free media were treated to HT-29 cells for 15 min and 30 min. Cells were washed twice with PBS and lysed with RIPA buffer (Biosolution, Korea) containing protease inhibitor and phosphatase inhibitor (Gendepot, USA) on ice. Lysed cells were collected using a scrapper and soluble lysate was obtained by centrifugation at 13,000 rpm for 15 min at 4°C. Total protein concentration in each sample was determined by the bicinchoninic acid (BCA) assay (Thermo, USA). Samples (20 μg) were loaded onto 8% and 12% PAGE to detect total STAT3 and phospho-STAT3 (pSTAT3), and β-actin, respectively. Mouse anti-STAT3 (1:1000, Cell Signaling Technology, USA), mouse anti-pSTAT3 (Tyr705; 1:2000, Cell Signaling Technology, USA), and rabbit anti-β-actin (1:5000, Proteintech, USA) antibodies were employed as a primary antibody. Subsequent reactions followed the same protocols as described above. For the β-actin detection, membrane was blocked with 5% skim milk (BD Difco, USA) in TBST for 20 min, and then protein band was developed through mouse anti-rabbit IgG-HRP (1:5000; Santa Cruz Biotechnology, USA).

### Cell Proliferation Assay

HT-29 cells were seeded into a 96-well plate (SPL, Korea) at a density of 1 × 10^4^ cells/well. After incubation for 24 hours, different concentrations of IL-22 (from 37.5 ng/ml to 600 ng/ml) were treated to each well for 48 hrs and 72 hrs. Cell proliferation was estimated using a WST-8 reagent (Biomax, Korea) which was incubated with cells for 30 min to 1 hour. The absorbance was measured at 450 nm with a microplate reader (BioTek, USA).

### Statistical Analysis

Data analysis was performed using Graphpad Prism 9 (San Diego, California, USA) software and Flowjo V10. The data were compared using a student’s t-test between the two groups and one-way analysis of variance (ANOVA) followed by post hoc tests (Tukey’s test) was used to compare more than two groups. Statistical significance is represented by *p < 0.05, **p < 0.01, ***p < 0.001.

## Results

### Bacterial Production of Recombinant Human Interleukin-22

Interleukin-22 (IL-22) which belongs to the class II cytokine family is described as a compact six-helix bundle (**Figure 1a**).[9, 11] The correct formation of two disulfide bonds between amino acids 40-132 and amino acids 89-178 is important to retain structural integrity and biological activity of IL-22.[9, 27] To promote the formation of native disulfide bridges, thioredoxin (Trx) was genetically fused with human IL-22, and the resulting gene was inserted into a pET32a bacterial expression vector (**Figure 1b** and **Figure S1**). We carried out bacterial expression of the recombinant Trx-tagged IL-22 (Trx-IL-22) in Origami B (DE3) *Escherichia coli* strain.[10, 28-30] As shown in **Figure 1c**, overexpression of Trx-IL-22 in bacteria was clearly observed in the protein band of after IPTG induction (AI) with a molecular weight of 35 kDa. Although most of expressed IL-22 aggregated in the form of inclusion body (IB), we successfully obtained about 4 mg of soluble Trx-IL-22 per liter culture. The collected elution fraction of Trx-IL-22 was subjected to western blot analysis to verify whether the purified protein is IL-22 by using anti-human IL-22 antibody (**Figure 1d**).[4, 31] In order to increase the purity of Trx-IL-22, size-exclusion chromatography (SEC) was subsequently carried out (**Figure 1e** and **Figure S2a**).[10] Unexpectedly, we observed that Trx-IL-22 eluted in two main peaks (A and B). Based on a calibration curve of SEC (**Figure S2b**), we identified that peak B with an estimated molecular weight of 42 kDa is a monomeric form of Trx-IL-22 while peak A is most likely a dimer (calculated as 106 kDa).[9, 10, 32] As previously reported, IL-22 is generally known to be functional as a monomer, but a dimeric form can be generated at a high protein concentration through hydrophobic and noncovalent interactions.[9, 33] Interestingly, we found that in the solution of peak A, insoluble aggregates were gradually formed in a time-dependent manner but this phenomenon did not occur in the peak B. The result suggests that the monomeric IL-22 is relatively stable compared to the dimeric form.

### Bioactivity Comparison of Monomeric and Dimeric Interleukin-22 *in vivo*

To compare the activity of dimeric and monomeric interleukin-22 (IL-22), we preliminary tested the treatment effect of either peak A or B of size-exclusion chromatography in a dextran sodium sulfate (DSS)-induced acute colitis mouse model.[26] The mice treated with 2.5% DSS lost their body weight by 20 % in day 9 and the colon length was shortened as a result of acute colitis compared to the control group (**Figure 2a** and **b**). Interestingly, mice injected with monomeric thioredoxin fused IL-22 (Trx-IL-22) of peak B showed notable protection from DSS-induced body weight loss and colon length shortening, which was not observed in dimeric Trx-IL-22-treated group (**Figure 2b** and **c**). The result suggests that the purified monomeric Trx-IL-22 have bioactivity to help repair and proliferation of injured colonic epithelial cells, whereas dimeric form of Trx-IL-22 was found to have no effect on ameliorating acute colitis in mouse model.[11, 14, 26, 34, 35] We speculate that the dimeric form exhibited low stability so that it could not play a protective role like the monomeric Trx-IL-22. Therefore, the monomeric Trx-IL-22 was selected for further downstream processes including an enzymatic reaction for Trx tag removal and a final polishing step using ion-exchange chromatography to evaluate the therapeutic potential of recombinant IL-22.

**Figure 2.**
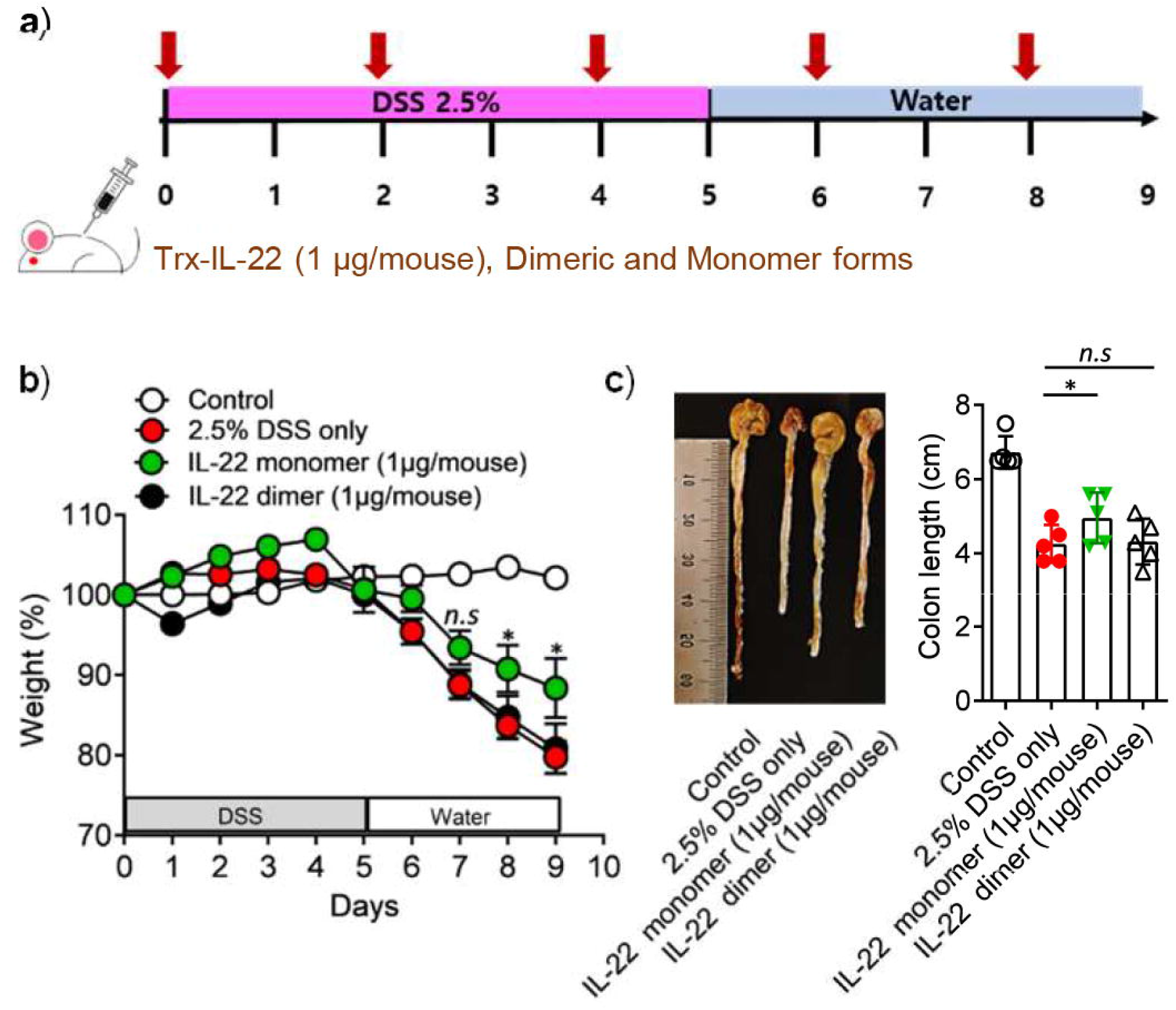
Biological effect by monomeric and dimeric form of interleukin-22 in DSS-induced colitis. **a)** Experimental scheme of Trx-IL-22 administration during 2.5% DSS-induced colitis progression. **b)** The daily percent change of body weight in each group. **c)** representative image and length of the colon. Data are expressed as mean ± SD, n=5 per group. *P < 0.05 (one-way ANOVA followed by Bonferroni test)

### Biochemical Characterization of Monomeric and Intact Interleukin-22

For the clinical application of recombinant proteins, it is inevitable to remove an unnecessary domain such as solubilizing tag and affinity tag from proteins of interest.[36] The downstream protein purification allows to prevent unwanted biological reactions and reduce the risk of adverse effects. To eliminate thioredoxin (Trx) tag from interleukin-22 (IL-22), enterokinase (EK) was treated to the monomeric Trx-IL-22 in which Trx tag is flanked by the recognition sequence of enterokinase. First, we determined the optimum incubation time of enterokinase (10 μg/ml) to allow an effective enzymatic reaction and avoid non-specific proteolysis. As a result, it was found that an incubation time of 2 hrs is sufficient to completely remove Trx tag from IL-22 by SDS-PAGE and western blot (**Figure 3a** and **Figure S3**).[31] After the enzyme reaction, monomeric IL-22 (mIL-22) was isolated through anion exchange chromatography with a linear salt-gradient owing to the remarkable difference in isoelectric point (pI) of IL-22 (6.8) and Trx (4.7). As shown in **Figure 3b**, mIL-22 was eluted in the first peak (A) with a remarkable high-purity (97.2%). To assess structural integrity of the resulting mIL-22, we performed circular dichroism (CD) spectroscopy, and revealed that the CD spectra of mIL-22 was very similar to that of the previously reported data showing α-helical structure (**Figure 3c**).[9, 37] Furthermore, computer prediction of protein secondary structure using CD data clearly provided that mIL-22 is mainly composed of alpha helices (90.8%). Based on these results, we demonstrated that human IL-22 can be effectively produced in bacteria as a monomeric and properly folded form.

**Figure 3.**
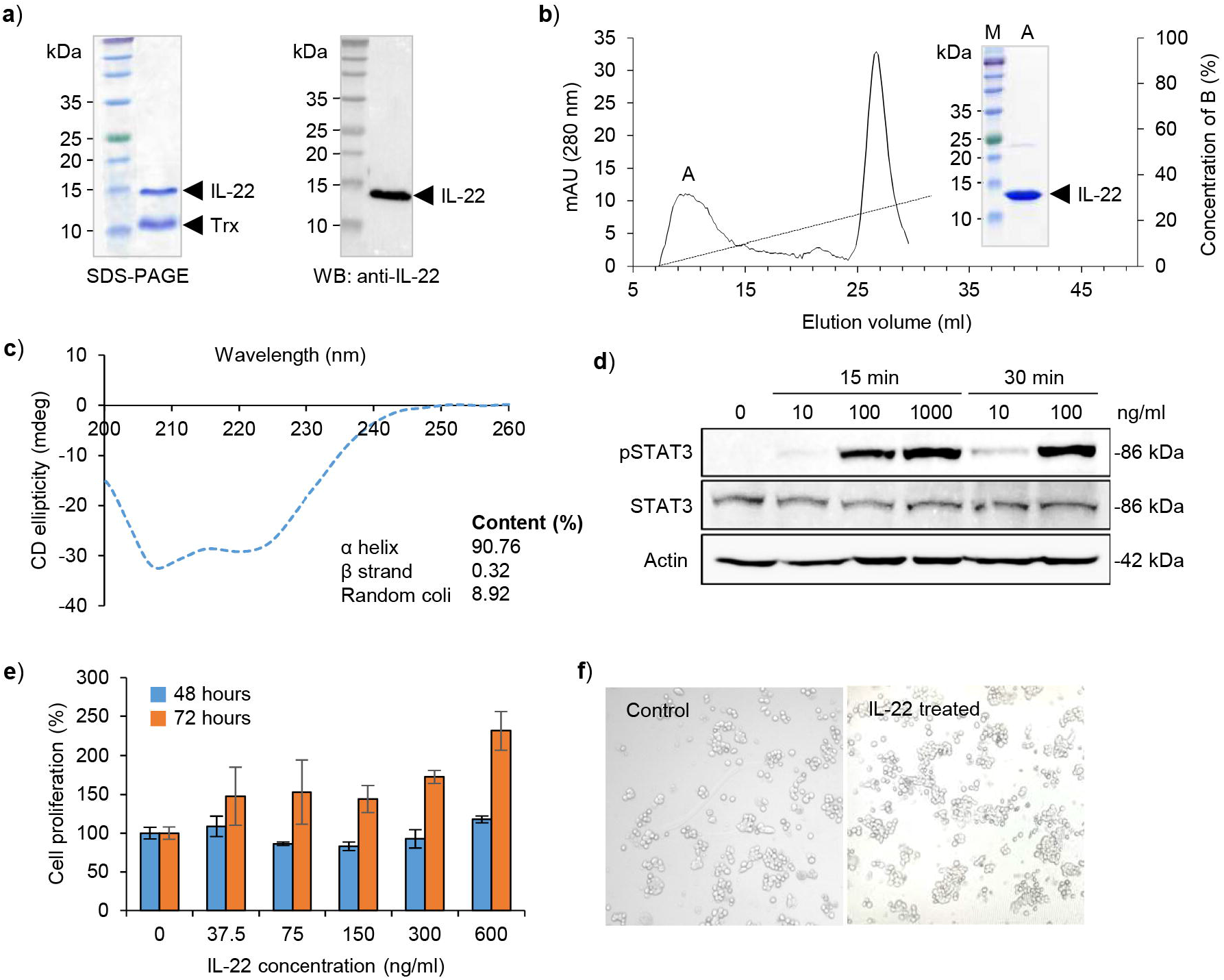
Characterization of monomeric interleukin-22 and evaluation of its biological activity. **a)** SDS-PAGE and western blot analysis of thioredoxin (Trx)-removed monomeric interleukin-22 (mIL-22) protein after enterokinase (EK) cleavage reaction. Enterokinase recognition sequence is located adjacent to the N-terminus of IL-22. For western blot analysis, anti-IL-22 antibody was used as a primary antibody. **b)** Anion exchange chromatography profile of purified mIL-22 protein from EK cleavage reaction using a linear NaCl gradient. Peak A containing mIL-22 was verified by SDS-PAGE analysis. **c)** Circular dichroism (CD) analysis of mIL-22. The result of secondary structure analysis was shown below the CD spectrum of IL-22. **d)** Western blot analysis of STAT-3 phosphorylation of HT-29 cells induced by mIL-22 treatment. STAT3, Phosphorylated STAT3 (pSTAT3), and actin were detected with increasing stimulation time and concentration of mIL-22. **e)** Dose-dependent effects of mIL-22 on proliferation of HT-29 cells. Graph showing the cell proliferation was presented depending on monomeric IL-22 treatment time (48 and 72 hours). **f)** Representative micrographs for verifying IL-22 promoted cell proliferation. HT-29 cells were cultured in serum free media with or without mIL-22 for 3 days. Bright-field images were obtained at 10x magnification.

Next, we investigated the biological activity of mIL-22 through cell-based assays. To elicit a biological response, IL-22 preferentially interacts with its cell surface receptors, IL-22 receptor 1 and IL-10 receptor 2.[11, 38] Formation of oligomeric IL-22/IL-22R1/IL-10R2 complexes triggers the activation of multiple cellular pathways, in particular signal transducer and activator of transcription 3 (STAT3) signaling.[35] To confirm the functionality of IL-22, we studied IL-22-induced STAT3 phosphorylation in HT-29 adenocarcinoma cell line overexpressing IL-22 receptors (IL-22R1 and IL-10R2).[4] Western blot analysis revealed that STAT3 phosphorylation (pSTAT3) was significantly occurred in time and concentration-dependent manner while there is no detectable band of pSTAT3 before IL-22 treatment (**Figure 3d**), indicating that recombinant mIL-22 is biologically active.[1, 31] Since STAT3 phosphorylation is closely related with the development and growth of cancer cells, we carried out cell viability assay to prove whether mIL-22 has the ability to stimulate cell proliferation.[39] As expected, mIL-22 treated cells showed in an approximately 2-fold increase in cell viability compared to the non-treated group (**Figure 3e** and **f**), implying that the functional mIL-22 has the potential as a biological agent for the study of related diseases, tissue regeneration and wound healing.[7]

### Investigation of the Ameliorative Effect of Interleukin-22 on DSS-induced Colitis

To assess the biological activity of monomeric interleukin-22 (IL-22), we treated mice with 1 μg of IL-22 every other day staring at the day of dextran sodium sulfate (DSS) administration for 5 times (**Figure 4a**). To induce colitis, mice were administered with 2.5 % DSS in the drinking water for 5 consecutive days, and it was replaced to fresh water at day 5. DSS treatment significantly decreased body weight (**Figure 4b**) and colon length of mice (**Figure 4c**), suggesting severe colitis developed in the colon.[26, 40] The levels of pro-inflammatory cytokines and chemokines including IL-1β, IL-6, IL-17A, C-X-C motif chemokine ligand 1 (CXCL1) and C-C motif chemokine ligand 2 (CCL2) were also highly elevated by DSS administration.[7] The intraperitoneal injection of IL-22 prevented body weight loss and reduced colon length shortening. In addition, the level of IL-6, IL-17A and CCL2 significantly reduced by IL-22 treatment (**Figure 4d**). In summary, we found that systemic injection of IL-22 attenuated DSS-induced colitis in murine model, suggesting that recombinant monomeric IL-22 could be a potent protein drug candidate to treat colitis.[41]

**Figure 4.**
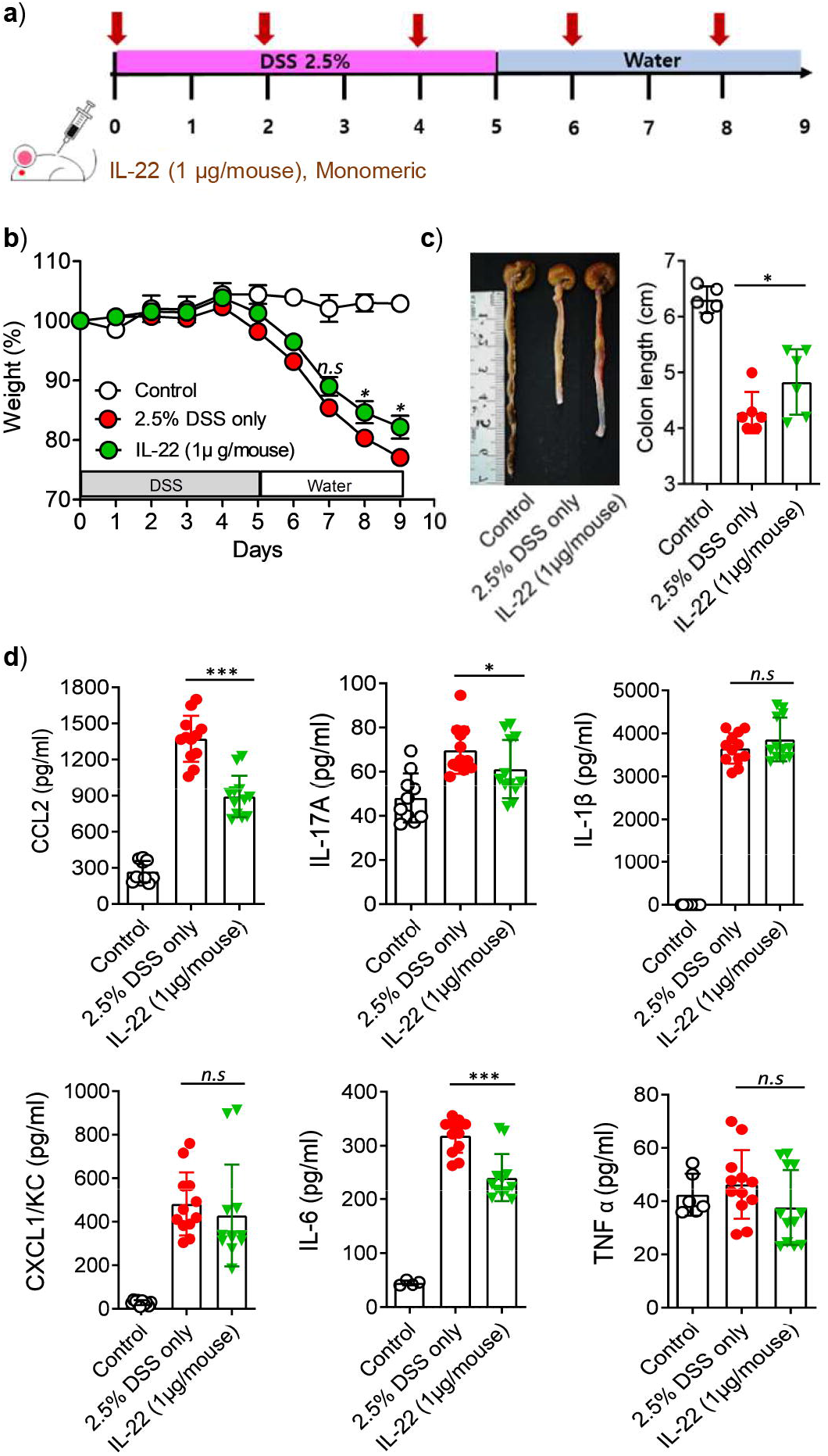
Monomeric interleukin-22 alleviates against colon damage in DSS-treated mice. **a)** The scheme of acute colitis along with monomeric IL-22 administration. **b)** The change of body weight percentage. **c)** Typical colon image and length. **d)** Inflammatory cytokines and chemokines were measured by ELISA in colon. Data are expressed as mean ± SD, n=5 per group. *P < 0.05, ***p <0.001 (one-way ANOVA followed by Bonferroni test)

## Discussion

Treatments for ulcerative colitis that have been used until recently include mesalazine, corticosteroids, monoclonal antibodies to TNF-α (infliximab, adalimumab), and immunosuppressants (cyclosporine, azathioprine).[42] Mesalazine is the first-line treatment for mild ulcerative colitis and is switched to other medications if the disease does not improve. However, since ulcerative colitis is difficult to cure, it is treated for the purpose of clinical maintenance and alleviation. In addition, if the condition is severe, a colectomy or immunosuppressant can be used, which increases the risk of infection or other disease as a side effect.[43] Therefore, it is necessary to develop a drug that is more effective and has fewer side effects than currently used drugs.

Interleukin-22 (IL-22) is a cytokine with high potential clinical relevance, and it is secreted from various immune cells including Th1, Th17, Th22, LTi and ILC3. IL-22 is a major regulatory cytokine for maintaining the homeostasis of epithelial cells in the intestine.[5, 7, 14, 15] In this regard, IL-22 regulates intestinal epithelial regeneration and intestinal barrier function by regulating junction protein expression and secretion of antimicrobial peptide and proteins including S100 and RegIII.[7, 16, 26] Previous studies have shown that IL-22 increases in inflammatory bowel disease (IBD) patients, but the disease was not dramatically improved. This may be due to the insufficient function of IL-22 by the inhibition of IL-22 binding protein (IL-22BP), a natural antagonist of IL-22, suggesting that extremely large amount of IL-22 may be required to overcome the IL-22BP-mediated inhibition.[7, 30, 38, 44] Indeed, several reports suggested that administration of IL-22 or IL-22Fc fusion protein exhibited sufficient protection against DSS-induced colitis in preclinical models.[15, 45, 46] On the contrary, however, blockade of IL-22 by neutralization antibody improved colitis induced by agonist anti-CD40 Ab or DNBS, suggesting complex role of IL-22 in intestinal inflammation and barrier function.[47] Recently, an IL-22Fc fusion protein, UTTR1147A, was assessed for its safety in Phase I clinical trial, and under evaluation for its efficacy in patients with ulcerative colitis, which can provide further information for the clinical applicability of IL-22.[36, 48]

In the current study, we presented that monomeric human IL-22 produced in bacteria has the therapeutic potential in attenuating interstitial inflammation in mouse models. IL-22, a multifunctional cytokine, is known to have a protective effect in various colitis models.[6, 13] In acute colitis models such as *Citrobacter rodentium*, IL-22 increased the expression of claudin-2, a tight junction protein, which leads to tissue repair and bacteria clearance.[47, 49-51] In addition, IL-22 inhibits prolonged inflammation to reduce damage within epithelial cells and directly targets LGR5+ stem cell to induce regeneration/proliferation of stem cell by STAT3 phosphorylation.[52, 53] Consequently, IL-22 contributes to protection of intestinal epithelial cells by inducing the regeneration of intestinal stem cell in acute tissue injury. On the contrary, however, high serum levels of IL-22 can lead to an aggravation of the disease and the chronic immune disorder in the intestines was regarded to be associated with IL-22-producing cells.[47, 54] Taken together, the role of IL-22 in the regulation of intestinal inflammation remains elusive, but IL-22 seems to be beneficial for homeostatic regulation of intestinal immunity and suppresses inflammation.[55]

Several reports suggested that IL-22 promoted tumor growth, especially in established tumors.[1, 31, 56, 57] This might be associated with the activation of STAT3 signaling pathway via IL-22-IL-22R interaction. On the contrary, however, IL-22 elicited protective role in early stage of tumor initiation.[7] Likewise, administration of anti-IL-22 Ab promoted tumor growth. Thus, although the role of IL-22 on colorectal cancer development is still controversial, it seems that IL-22 can act to tumor initiation.

Given the pleiotropic action of IL-22, understanding the underlying mechanisms of IL-22 in disease is of fundamental importance for maximizing therapeutic benefits and providing insight into potential therapeutic implications. Securing sufficient amounts of bioactive IL-22 protein is a prerequisite for facilitating the evaluation of biochemical and biological properties of IL-22. We thus genetically engineered human IL-22 through codon optimization and thioredoxin fusion to improve the translation efficiency and accelerate disulfide bond formation, leading to high-level soluble expression of IL-22 in bacteria. Interestingly, the expressed IL-22 exhibited in the form of a biologically active monomer and a non-functional homodimer. We found clearly that monomeric IL-22 (mIL-22) possesses the ability to attenuate DSS-induced acute colitis, whereas dimeric IL-22 (dIL-22) is prone to aggregation without observable biological activity *in vivo*. However, the therapeutic efficacy of mIL-22 in colitis seems to be insufficient compared to other chemical reagents, so genetic engineering of IL-22 is required to generate more potent variants. As mentioned above, IL-22 can tightly interact with two endogenous proteins, IL-22R1 and IL-22BP in a competitive manner. IL-22/STAT3 signaling is triggered by the complex formation between IL-22 and IL-22R1, but conversely, the signaling pathway is also strongly blocked by IL-22BP, a soluble decoy receptor.[38] Accordingly, modulating the binding affinity of IL-22 to IL-22R1 to maximize the agonistic effect of IL-22 can be considered a promising strategy.[30] To this end, we plan to conduct further studies to enhance the binding affinity of IL-22 to IL-22R1 and to modify critical residues of IL-22 in IL-22BP interactions through directed evolution and computational protein design.

In conclusion, we demonstrated that mIL-22 can be effectively produced using bacterial expression system and ameliorate DSS-induced colitis in mouse models by inhibiting the expression of pro-inflammatory cytokines. Considering the increasing number of patients with inflammatory disorders of the gastrointestinal tract, it is anticipated that the recombinant mIL-22 can be developed as a biological agent with promising therapeutic potential in colitis and other related diseases.

## Acknowledgements

This work was supported by the National Research Foundation (NRF) of Korea (2019R1I1A3A01047208, 2019R1I1A1A01058773, 2020R1A2B5B02001552 and 2020R1A5A8019180), the National Research Facilities & Equipment Center (2019R1A6C1010006), and Kangwon National University (520170494).

## Author Contributions

Conceptualization: Lee JJ, Ko HJ; Data curation: Hong EH, Ryu Y; Funding acquisition: Hong EH, Ryu Y, Lee JJ, Ko HJ; Investigation: Kim S, Hong EH, Lee CK; Methodology: Ryu Y, Jeong HJ, Heo S; Supervision: Lee JJ, Ko HJ; Writing - original draft: Kim S, Hong HE, Lee CK, Ryu Y; Writing - review & editing: Lee JJ, Ko HJ.

