## Supplemental Information for "Amelioration of DSS-induced Acute Colitis in Mice by Recombinant Monomeric Human Interleukin-22"

[1]Department of Biochemistry, Kangwon National University, Chuncheon 24341, South Korea. [2]Laboratory of Microbiology and Immunology, Department of Pharmacy, Kangwon National University, Chuncheon 24341, South Korea. [3]Institute of Life Sciences (ILS), Kangwon National University, Chuncheon 24341, South Korea. [4]Global/Gangwon Innovative Biologics-Regional Leading Research Center (GIB-RLRC), Kangwon National University, Chuncheon 24341, South Korea. [5]These authors contributed equally to this work.

**>Thioredoxin-tagged IL-22 (Trx-IL-22)**

MSDKIIHLTDDSFDTDVLKADGAILVDFWAEWCGPCKMIAPILDEIADEYQGKLTV
AKLNIDQNPGTAPKYGIRGIPTLLLFKNGEVAATKVGALSKGQLKEFLDANLAGSG
SGHMHHHHHHSSGLVPRGSGMKETAAAKFERQHMDSPDLGTDDDDKAMGAPISS
HCRLDKSNFQQPYITNRTFMLAKEASLADNNTDVRLIGEKLFHGVSMSERCYLMK
QVLNFTLEEVLFPQSDRFQPYMQEVVPFLARLSNRLSTCHIEGDDLHIQRNVQKLK
DTVKKLGESGEIKAIGELDLLFMSLRNACI

**Figure S1. Complete amino acid sequences of thioredoxin-tagged human interleukin-22 constructed in this study.** Interleukin-22, thioredoxin and histidine tag are colored in green, blue and yellow, respectively. Enterokinase recognition sequence is highlighted in red color.

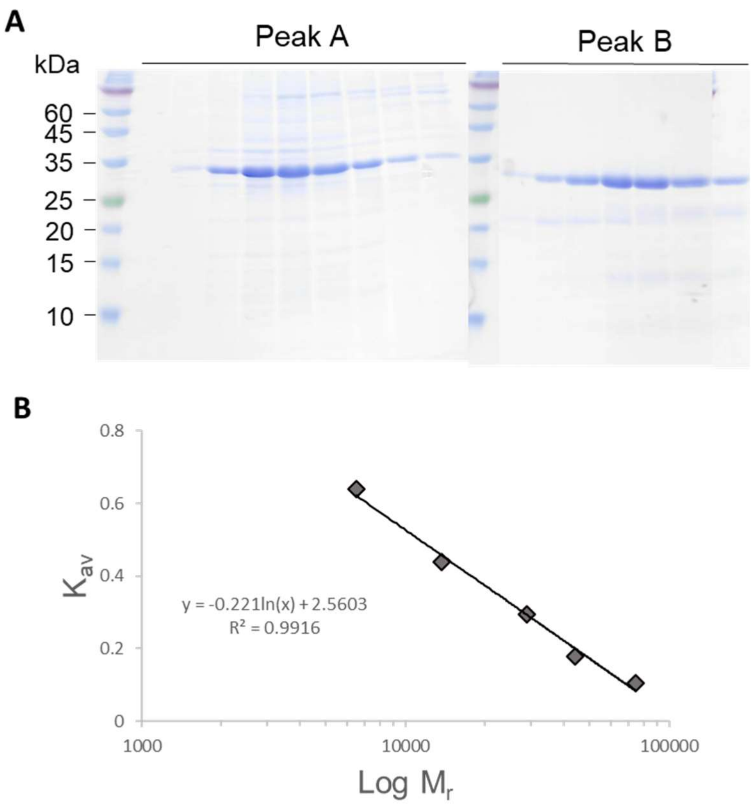


**Figure S2. Purification of thioredoxin-tagged interleukin-22 (Trx-IL-22) by size exclusion chromatography. a**) All elution fractions of peak A and B containing Trx-IL-22 shown in **Figure 1e** were analyzed by SDS-PAGE. **b**) Calibration curve of HiLoad 16/60 Superdex 75 column by standard molecular weight proteins (Conalbumin: 75 kDa, Ovalbumin: 43 kDa, Carbonic Anhydrase: 29 kDa, Ribonuclease A: 13.7 kDa, Aprotinin: 6.5 kDa).

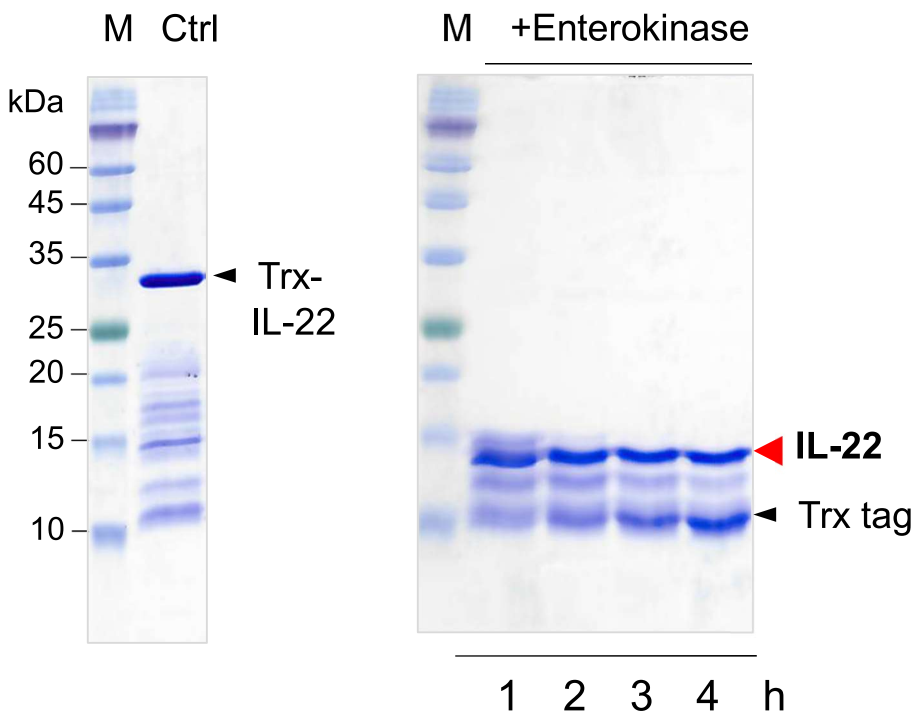


**Figure S3. SDS-PAGE analysis of cleaved monomeric interleukin-22 depending on enterokinase cleavage reaction time.** The left SDS-PAGE showed monomeric thioredoxin-tagged interleukin-22 (Trx-IL-22) before enterokinase treatment. During the enterokinase cleavage reaction (1, 2, 3 and 4 hours), monomeric IL-22 was found as an independent form without Trx tag because of the specific cleavage at DDDDK site located between Trx tag and IL-22 (right SDS-PAGE).
